# *In Vitro* Analysis of the Anti-viral Potential of nasal spray constituents against SARS-CoV-2

**DOI:** 10.1101/2020.12.02.408575

**Authors:** Mark L Cannon, Jonna B. Westover, Reiner Bleher, Marcos A. Sanchez-Gonzalez, Gustavo Ferrer

**Affiliations:** Department of Otolaryngology Division of Dentistry, Feinberg School of Medicine, Long Grove IL USA; Ann and Robert Lurie Children’s Hospital of Chicago; Chicago, IL, USA; Institute for Antiviral Research, Utah State University, Logan, Utah, USA; Department of Materials Science & Engineering, Northwestern University, Evanston, IL, USA; Lake Erie College of Osteopathic Medicine, Bradenton, FL, USA; Medical Innovations, Nova Southeastern University, Davie, FL, USA; Research & Development, Aventura Pulmonary Institute, Miami, FL, USA

## Abstract

Viral pandemics have taken a significant toll on humanity and the world now is contending with the SARS-CoV-2 epidemic. Readily available economical preventive measures should be immediately explored. Xylitol has been reported to reduce the severity of viral infections as well as the severity of pneumonia, and increase the survivability of animal subjects. Since pneumonia and acute respiratory distress syndrome are potentially fatal complications of COVID-19, the present study tested the *in vitro* effectiveness of xylitol against SARS-CoV-2. Virus titers and LRV of SARS-CoV-2, were incubated with a single concentration of nasal spray. Toxicity was observed in the top dilution (1/10). Virus was seen below that dilution so it did not affect calculations of virus titer or LRV. After a 25-minute contact time, the nasal spray (11% Pure Xylitol, 0.85%NaCL (Saline), and 0.20% grapefruit seed extract) reduced virus from 4.2 to 1.7 log10 CCID50 per 0.1 mL, a statistically significant reduction (P<0.001) of 2.5 log10 CCID50. STEM Images obtained at the BIoCryo Laboratory revealed virus contained on the cell wall but none intra-cellular, possibly due to D-xylose (xylitol) production of glycoaminoglycans decoy targets. Xylitol and grapefruit seed extract are not exotic nor expensive rare high technology answers to viral epidemics. The potential in saving lives and the economies of the world by using X-GSE combination therapy should inspire large clinical trials, especially in those nations whereas the healthcare system would be dangerously compromised by the adoption of less effective and significantly more financially demanding therapies. Because there are no risk factors in using the X/GSE combination therapy, and the nasal spray is over the counter available without a prescription, and the spray allows for comfortable long term mask-wearing, adoption of this preventive anti-viral therapy should be encouraged.

## Introduction

Viral pandemics have taken a significant toll on humanity and the world now is contending with the SARS-CoV-2 epidemic. As of November, 28^st^, 2020, the estimate of cases and related fatalities for the world are reported as 61,877,685 and 1,447,246 respectively.^1^ The societal cost of COVID-19 is very difficult to measure, but millions have lost their livelihoods and the Federal (USA) expenditures top 3 trillion dollars, more than the amount spent on all scientific research in the history of federal expenditures.^2^ And yet there is little to show for this Herculean effort and expenses as of this date. For example, the total cost of World war II was 4 trillion dollars in today’s dollars over 4 years with 481,000 fatalities for the United States of America. It appears that more Americans will die early in just one year due to the COVID-19 epidemic. Readily available and economical preventive measures should be immediately explored.

Newly published research has demonstrated the antiviral properties of polyols. Xylitol has been reported to reduce the severity of viral infections. The effect of dietary xylitol on hRSV infection was investigated in a mouse model with significant results reported.^4^ The mice received xylitol for 14 days before virus exposure and for a further three days post-viral exposure. The mice receiving xylitol had significantly reduced viral lung titers than the controls receiving phosphate-buffered saline (PBS). Fewer CD3+ and CD3+CD8+ lymphocytes, whose numbers are indicative of inflammatory status, were recruited in the mice receiving xylitol. These results demonstrated improved hRSV infection outcomes and reduced inflammation-associated immune responses to hRSV infection with dietary xylitol. The same researchers previously reported positive effects of xylitol on mice with influenza A virus infection (H1N1) also with a decrease in recruitment of inflammatory lymphocytes.^5^ It has been reported that a decrease in CD3+CD8+ lymphocytes is a predictor of mortality for COVID-19 patients. The antiinflammatory and antiviral properties of D-xylose/xylitol in respiratory conditions are subject to a patent application (number WO1999048361A1) filed in 1998 in the United States. 6 Subsequently, xylitol is a main active ingredient in nasal spray products, such as Xlear Sinus Care.

Xylitol has also been demonstrated to reduce the severity of pneumonia, and increase the survivability of animal subjects.^7, 8^ Pneumonia and acute respiratory distress syndrome are potentially fatal complications of COVID-19. 9 Interestingly, xylitol has been used in clinical trials to decrease *Pseudomonas aeruginosa* in patients with cystic fibrosis.^10, 11^ The idea of using xylitol in the ventilators dedicated to COVID-19 patients was not, however, put into practice due to the number of clinical trials with other funded treatments that later demonstrated limited success. ^12^ Further research into the effectiveness of xylitol against SARS-CoV-2 is therefore required.

## Materials and Methods

### Virucudal Assay

SARS-CoV-2, USA-WA1/2020 strain, virus stock was prepared before testing by growing in Vero 76 cells. Culture media for prepared stock (test media) was MEM with 2% fetal bovine serum and 50 μg/mL gentamicin. The compound was mixed directly with virus solution so that the final concentration was 90% of the compound preparation and 10% virus solution. A single concentration was tested in triplicate. Test media without virus was added to one tube of the prepared compound to serve as toxicity controls. Ethanol (70%) was tested in parallel as a positive control and water only as a virus control. Solution and virus were incubated at room temperature (22 ± 2°C) for 25 minutes. The solution was then neutralized by a 1/10 dilution in test media.

Surviving virus from each sample was quantified by standard end-point dilution assay. Briefly, samples were serially diluted 1/10 in test medium. Then 100 μL of each dilution were plated into quadruplicate wells of 96-well plates containing 80-90% confluent Vero 76 cells. Plates were incubated at 37 ± 2°C with 5% CO2 for 6 days. Each well was then scored for the presence or absence of virus. The end-point titers (CCID50) values were calculated using the Reed-Muench (1948) equation. Three independent replicates of each sample were tested, and the average and standard deviation were calculated. Results were compared with untreated controls by one-way ANOVA with Dunnett’s multiple comparison tests using GraphPad Prism (version 8) software. Controls: Virus controls were tested in water and the reduction of virus in test wells compared to virus controls was calculated as the log reduction value (LRV). Toxicity controls were tested with media not containing virus to determine if the samples were toxic to cells. Neutralization controls were tested to ensure that virus inactivation did not continue after the specified contact time, and that residual sample in the titer assay plates did not inhibit growth and detection of surviving virus. This was done by adding toxicity samples to titer test plates then spiking each well with a low amount of virus that would produce an observable amount of CPE during the incubation period.

### Electron Microscopy

Cell pellets were fixed in 3% glutaraldehyde, 2% formaldehyde in 0.1 M PIPES, pH 7.2 for 72 hours. After fixation, cells were enrobed in 10% gelatin, rinsed in 0.1 M PIPES for 3 x 10 minutes and postfixed in 1% osmium tetroxide for 1 hour. After two rinses for 10 minutes in DI water, cells were en bloc stained with 2% uranyl acetate in DI water and rinsed 2 x 5 minutes with DI water. Samples were dehydrated in an ascending series of ethanol (25%, 50%, 75%, 95% and 3 x 100%) for 15 minutes each and infiltrated at RT with EMBed 812 resin/ethanol mixture 1:1 for 30 minutes, 3:1 for 60 minutes, and with pure resin overnight. The next day, samples were transferred into fresh resin in silicon molds and polymerized at 65□C for 48 hours. Sections of ca. 80 nm thickness were generated with a diamond knife (Diatome, Hatfield, PA) using a Leica Ultracut-S ultramicrotome. The sections were placed on TEM grids and images were recorded with a Hitachi HD2300 STEM at 200 kV acceleration voltage.

(Chemicals were from EMS, Hatfield, PA)

|  |  |  |  |
| --- | --- | --- | --- |
| Glutaraldehyde 25% | EMS | 1x10 mL | #16200 |
| Formaldehyde 16% | EMS | 1x10 mL | #15700 |
| PIPES buffer | EMS | 450 mL | #11610 |
| EMBed812 | EMS | Kit | #14750 |
| Grids: | EMS |  | #G200-Cu |

## Results

### Virucidal Assay

Virus titers and LRV of SARS-CoV-2, when incubated with a single concentration of nasal spray, are shown in Table 1. Toxicity was observed in the top dilution (1/10). Virus was seen below that dilution so it did not affect calculations of virus titer or LRV. After a 25-minute contact time, the nasal spray reduced virus from 4.2 to 1.7 log10 CCID50 per 0.1 mL, a statistically significant reduction of 2.5 log10 CCID50. Neutralization controls demonstrated that residual sample did not inhibit virus growth and detection in the endpoint titer assays. Virus controls and positive controls performed as expected.

**Table 1.** Virucidal efficacy of nasal spray against SARS-CoV-2 after a 25-minute incubation with virus at 22 ± 2°C.

|  | Tested<br>Concentration | Incubation<br>Time | Virus<br>Titer<sup>a</sup> | LRV<sup>b</sup> |
| --- | --- | --- | --- | --- |
| Xlear Nasal Spray | 90% | 25-minute | <0.67 | 3.33 |
| Xlear Rescue Spray | 90% | 25-minute | <0.67 | 3.33 |
| XRS +50 ppm silver | 90% | 25-minute | <0.67 | 3.33 |
| XyChlor Spray, pH 6.1 | 90% | 25-minute | <0.67 | 3.33 |
| Chlorhexidine 0.03% | 90% | 25-minute | 3.67 | 0.33 |
| Chlorhexidine 0.015% | 90% | 25-minute | 3.67 | 0.33 |
| Ethanol | 67.5% | 25-minute | <0.67 | 3.33 |
| Virus Control | na | 25-minute | 4.0 | na |

Concentration Incubation Virus Titera LRVb
Nasal spray 90% 25-minute $1.7 \pm 0.0$ 2.5\*\*\*
Ethanol 67.5% 25-minute $1.0 \pm 0.6$ 3.2\*\*\*
Virus Control na 25-minute $4.2 \pm 0.4$ na
a Log10 CCID50 of virus per 0.1 mL, average of 3 replicates ± standard deviation
b LRV (log reduction value) is the reduction of virus compared to the virus control
\*\*\*P < 0.001 by one-way ANOVA and Dunnett's post-test compared with untreated virus control (water). For wells with undetectable virus a value equal to the lower limit of detection was assigned for statistical analyses.

### Imaging

Scanning transmission electron microscope Images obtained at the BIoCryo Laboratory (Northwestern University) revealed virus contained on the cell wall but none intra-cellular, possibly due to D-xylose (xylitol) production of glycoaminoglycans decoy targets. A very recently published article correlated d-xylose and xylitol to the severity and morbidity related factors in COVID-19. 13 D-xylose is the initiating element for sulfated glycosaminoglycans (GAG) that attach to the core protein. D-xylose can be derived from xylitol by d-xylose reductase action replenishing this carbohydrate that is targeted by the SARS-CoV-2 virus. If the virus attaches to the d-xylose position on the GAG, such as heparin sulfate, the virus can then contact the ACE2 receptor. Additionally, xylitol serves as a decoy target for the virus, preventing it from successfully reaching the ACE2 receptor. Further research into the mechanism that xylitol initiates with cells to prevent viral penetration and replication is essential and should be prioritized.

## Discussion

Xylitol has a long history of being safe and beneficial in preventing bacterial pathogen infections.^14^ It is considered a prebiotic due to its positive effect on the microbiome, reducing pathogenic proliferation.^15^ The use of xylitol in oral health to prevent dental caries and periodontal disease has been well documented as safe and effective.^16, 17^ Studies have shown xylitol inhibits the formation of mixed species biofilms, which include *Porphyromonas gingivalis* in vitro.^16, 18^ Long term clinical studies have demonstrated that children with dental problems grow up to be adults with heart disease. For example, Kids with dental abscesses were followed for 27 years and they developed pre-clinical signs of coronary heart disease. 19 In addition, general morbidity and mortality rates are very closely associated with advanced periodontal disease, and there are also well documented connections to inflammatory Alzheimer’s disease and atherosclerosis.^20–25^

By inhibiting P gingivalis with xylitol and erythritol, the innate and adaptive immune response of the human host should be more robust.^26^ Also, the possibility of salivary spread of oral pathogens should be reduced, preventing onset of the acute respiratory distress syndrome. Indeed, there have been three important recently published peer review articles, two in Medical Hypothesis and one in the British Dental Journal that reinforce how important oral health care is in regards to COVID-19 itself. Periodontal disease is another of the pre-disposing co-morbidities.^27^ This is certainly not surprising as PD creates systemic inflammation, increase in proinflammatory cytokine levels. This would exacerbate the cytokine storm of COVID-19, and the oral pathogens in the saliva could cause an increase in the pneumonia risk.^28^

Xylitol and grapefruit seed extract are not exotic nor expensive rare high technology answers to viral epidemics. The potential in saving lives and the economies of the world by using X-GSE combination therapy should inspire large clinical trials, especially in those nations whereas the healthcare system would be dangerously compromised by the adoption of less effective and significantly more financially demanding therapies. Also, the potential mechanism by which xylitol may pose as a decoy target for the SARS-CoV-2 virus was just recently published. ^13^ Because there are no risk factors in using the X/GSE combination therapy, and the nasal spray is over the counter available without prescription, and the spray allows for comfortable long term mask wearing, adoption of this preventive anti-viral therapy should be encouraged.

## Conclusion

Combination therapy with GSE and xylitol may prevent spread of viral respiratory infections not just for SAR-CoV-2 but also for future H1N1 or other viral epidemics. GSE significantly reduces the viral load while xylitol prevents the virus attachment to the core protein on the cell wall.

**Figure 1.**
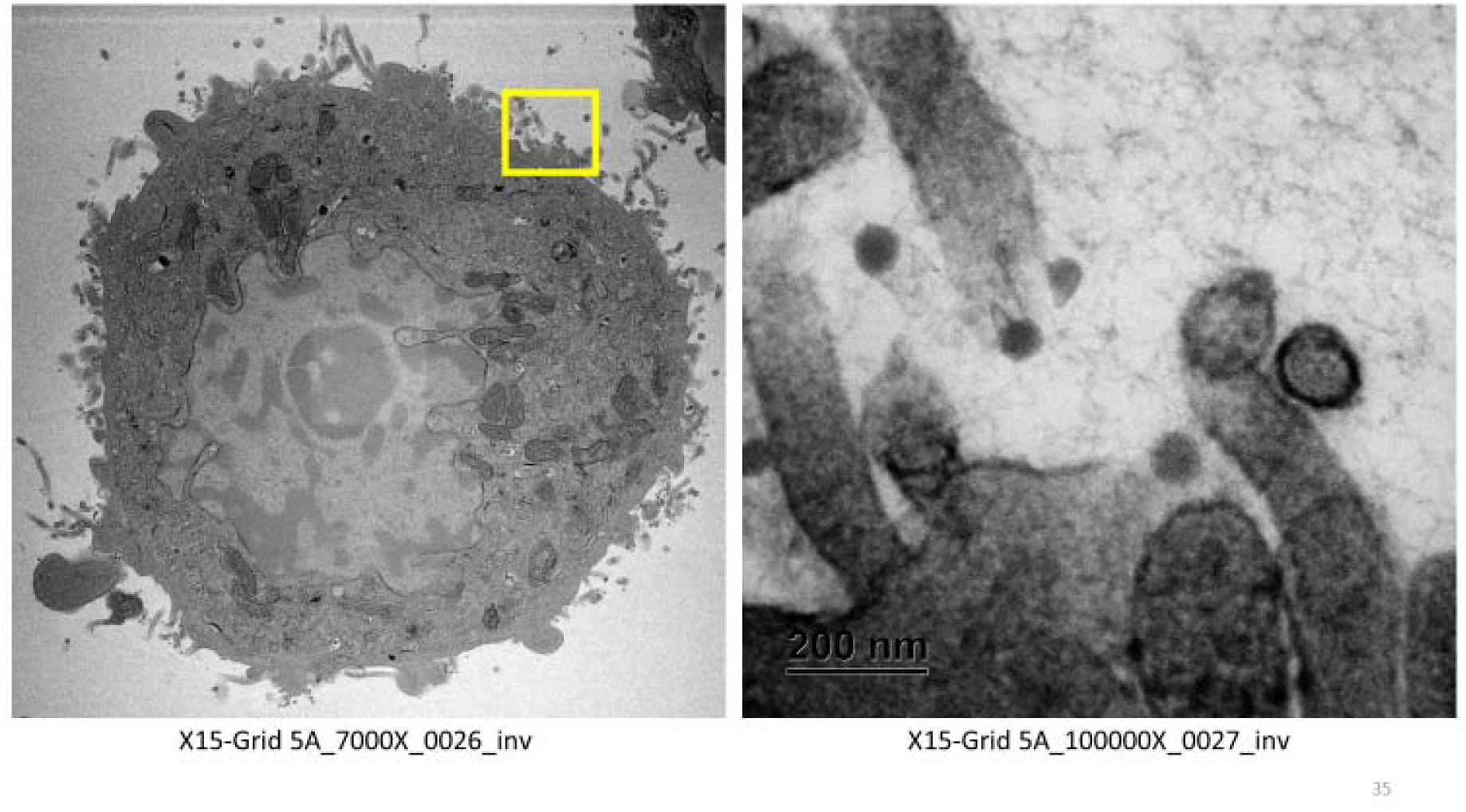
SARS-CoV-2 virus outside Vero 76 immortalized cells.

**Figure 2.**
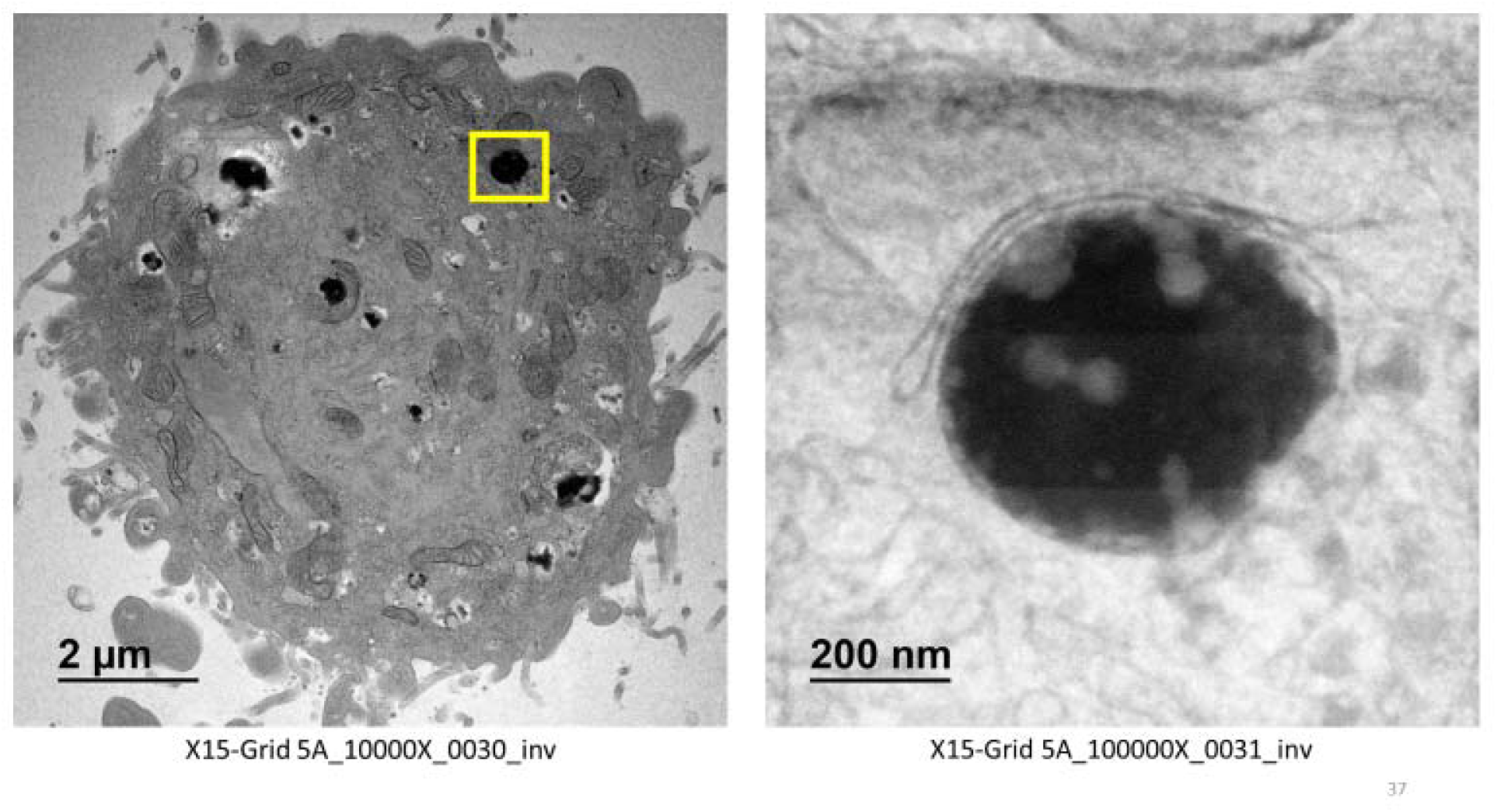
SARS-CoV-2 on cell wall. No viral inclusion bodies noted within cell cytoplasm.

## Acknowledgement

“This work made use of the BioCryo facility of Northwestern University’s NUANCE Center, which has received support from the SHyNE Resource (NSF ECCS-2025633), the IIN, and Northwestern’s MRSEC program (NSF DMR-1720139).”

